# Dilated Virchow-Robin Spaces are a Marker for Arterial Disease in Multiple Sclerosis

**DOI:** 10.1101/2023.02.24.529871

**Authors:** Benjamin V. Ineichen, Carmen Cananau, Michael Plattén, Russell Ouellette, Thomas Moridi, Katrin B. M. Frauenknecht, Serhat V. Okar, Zsolt Kulcsar, Ingrid Kockum, Fredrik Piehl, Daniel S. Reich, Tobias Granberg

## Abstract

Virchow-Robin spaces (VRS) have been associated with neurodegeneration and neuroinflammation. However, it remains uncertain to what degree non-dilated or dilated VRS reflect specific features of neuroinflammatory pathology. Thus, we aimed at investigating the clinical relevance of VRS as imaging biomarker in multiple sclerosis (MS) and to correlate VRS to their histopathologic signature. In a cohort study comprising 205 MS patients (including a validation cohort) and 30 control subjects, we assessed the association of non-dilated and dilated VRS to clinical and magnetic resonance imaging (MRI) outcomes. Brain blocks from 6 MS patients and 3 non-MS controls were histopathologically processed to correlate VRS to their tissue substrate. The count of dilated centrum semiovale VRS was associated with increased T1 and T2 lesion volumes. There was no systematic spatial colocalization of dilated VRS with MS lesions. At tissue level, VRS mostly corresponded to arteries and were not associated with MS pathological hallmarks. Interestingly, dilated VRS in MS were associated with signs of small vessel disease. Contrary to prior beliefs, these observations suggest that VRS in MS do not associate with accumulation of immune cells. But instead, these findings indicate vascular pathology as a driver and/or consequence of neuroinflammatory pathology for this imaging feature.

## Introduction

Perivascular spaces surround blood vessels in the brain^1^. Macroscopically visible perivascular spaces, also referred to as Virchow-Robin spaces (VRS) or enlarged perivascular spaces, are readily detectable by magnetic resonance imaging (MRI)^2,3^. VRS appear as thin linear or small punctate structures with a signal similar to CSF on MRI^4^. Normally, VRS are smaller than 2 mm in diameter; yet, due to unknown reasons, VRS may dilate and present with diameters ≥ 2 mm^5^.

Although there is an ongoing debate about the clinical relevance of VRS^6,7^, a large body of evidence indicates their association with ageing, vascular risk factors like hypertension^8^, vascular diseases such as small vessel disease^9^ or stroke^10,11^ as well as neurodegenerative diseases^12^.

Accumulating evidence also suggests an association of VRS with neuroinflammatory disorders, including multiple sclerosis (MS)^3^. MRI studies have shown that MS patients harbor a higher VRS burden compared to control subjects^13,14^. This has been confirmed in a recent meta-analysis^15^. Furthermore, albeit not reproduced so far, increase of overall VRS volume may precede the emergence of contrast-enhancing MS lesions^16^. Finally, it has been shown that MS patients have elevated numbers of dilated VRS^17^.

Together, the exact roles of both nondilated and dilated VRS in MS pathogenesis are insufficiently understood. To further corroborate nondilated and/or dilated VRS as an imaging biomarker in MS, several open questions need to be answered: (1) Are VRS cross-sectionally or longitudinally associated with clinical or imaging outcomes in MS? (2) What is the topographical distribution of VRS, and do they coincide with MS lesions? (3) What is the longitudinal evolution of VRS in MS? And (4) What is the anatomical and/or pathological substrate of VRS in MS, and could they also be associated with signs of vascular diseases in MS?

Here, we studied nondilated and dilated VRS in the largest MS patient cohort to date, collectively comprising 205 patients (including 24 patients with longitudinal MRI scans) and 30 controls, with prospective collection of clinical outcomes and standardized T1- and T2-weighted (T1w, T2w) MRI. Finally, in brain samples from MS donors and non-MS controls with corresponding pre- and postmortem MRI scans, we assessed the anatomical and pathological signature of VRS on the tissue level.

## Materials and Methods

### Data availability

The Anonymized data are available from the corresponding author upon reasonable request and in accordance with current legislation.

### Study design and subjects

Our exploratory cohort comprised 142 MS patients and was derived from the *Stockholm Prospective Assessment of MS* (STOP-MS) from the Karolinska University Hospital in Sweden (including 24 patients with longitudinal data). The validation cohort constitutes a subcohort of the *MultipleMS* and *MyelinMS* studies conducted at the same institution and comprised 63 MS patients. The control cohort consisted of 30 sex- and age-matched subjects scanned on the same scanner. All studies include longitudinal follow-up including standardized MRI scanning protocols at 3 Tesla and longitudinal clinical evaluations. Inclusion criteria for our study: subjects with T1w and T2w as well as T2w-fluid-attenuated inversion recovery (FLAIR) MRI. For the longitudinal cohort, patients with relapses during time of imaging were excluded. In addition, for the histopathologic assessment, brains from 9 subjects were included (6 MS patients, 3 non-MS patients). Patients with a history of stroke or cardial infarction were excluded.

### Clinical outcomes

Clinical disability of the patients was assessed using the Expanded Disability Status Scale (EDSS)^18^. Cognitive functioning was measured by the Symbol Digit Modalities Test (SDMT)^19^. The cognitive scores were normalized to age- and sex-adjusted *z*-scores based on normative data^20^. For the STOP-MS cohort, vascular risk profiles were compiled based on the presence of diabetes, dyslipidemia, hypertension, smoking status, and antiplatelet therapy.

### *In vivo* MRI

Imaging was performed on the same Siemens Trio/PRISMA^Fit^ 3 T MRI scanner (Siemens Healthcare, Erlangen, Germany) with a 20-channel head coil. The protocol included: (1) A 3D T1w sequence (repetition time = 1900 ms, echo time = 3.39 ms, inversion time = 900 ms, flip angle 9 degrees, spatial resolution 1.0×1.0×1.5 mm^3^), for VRS detection, atrophy measurements, and T1 lesion detection; (2) The same 3D T1w sequence was applied after intravenous administration of a standard dose (0.2 ml/kg) of gadoteric acid (Dotarem) after a minimum of 5 minute delay, which was administered as part of the clinical routine; (3) A 3D T2w sequence (repetition time = 3200 ms, echo time = 388 ms, flip angle = 120 degrees, spatial resolution 1.0×1.0×1.0 mm^3^); and (4) A 3D T2w FLAIR (TR = 6000 ms, TE = 388 ms, TI = 2100 ms, flip angle 120 degrees, spatial resolution 1.0×1.0×1.0 mm^3^) for T2 lesion detection.

### Lesion and brain atrophy measures

FreeSurfer version 6.0.0 was used to perform automated cross-sectional and longitudinal brain volume measures and segment T1 hypointense lesions^21^. For volumetric analyses, gray and white matter volume as well as whole brain volume were normalized to the estimated total intracranial volume, resulting in the corresponding tissue fractions^22^. To estimate longitudinal brain atrophy rates, the longitudinal stream of FreeSurfer was used to obtain the white matter, gray matter, and brain parenchymal fractions (WMF, GMF, and BPF). Numerically, the latest available MRI measurement was subtracted by the earliest available MRI measurement, divided by the time in between the scans. Lesion Segmentation Toolbox 2.0.15^23^ for SPM12 was used to perform cross-sectional and longitudinal T2 lesion volume segmentations based on the FLAIR volumes. The volumes of contrast-enhancing lesions were manually segmented on the post-contrast 3D T1-weighted images using ITK-SNAP^24^.

### Assessment of VRS

VRS were defined according to the STRIVE criteria, i.e., as fluid-filled spaces that follow the typical course of a vessel through gray or white matter and with similar signal intensity to CSF on all sequences^1^. VRS were quantified on the axial and coronal reformatted T1-weighted images by a resident in neuroradiology (BVI) with 2 years of training using the open source DICOM viewer Horos. FLAIR images were consulted to differentiate VRS from MS lesions (hyperintense on T2 FLAIR) or lacunar infarcts (commonly with hyperintense rim). Following prior work^13^, four brain levels were used as landmarks to assess VRS: (1) the hand knob at the high convexity/centrum semiovale, (2) at the widest part of the crus anterius, (3) at the level of the anterior commissure and (4) in the midbrain at the largest interpeduncular distance (**Figure 1A-E**). We decided post-hoc to pool VRS at the two levels of the basal ganglia by using their mean to reduce the number of statistical tests. Additionally, VRS outcomes were rated by an experienced neuroradiologist (CC) on 25 randomly selected MRI scans to estimate the inter-rater agreement.

**Figure 1:**
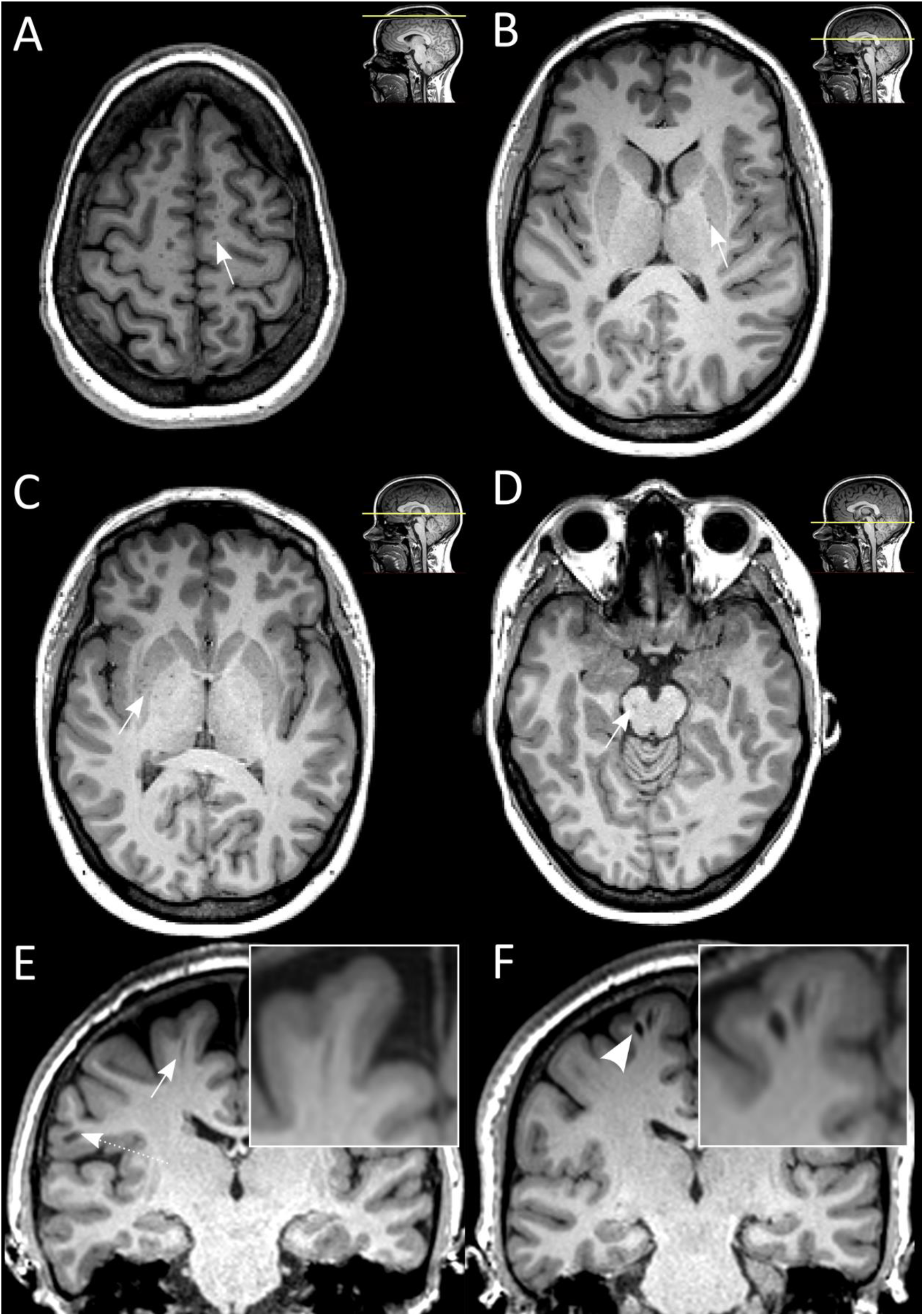
Virchow-Robin spaces (VRS) are readily detectable on T1-weighted brain MRI scans. <u>A-D</u>: VRS were assessed at 4 different brain levels: centrum semiovale (A), basal ganglia, crus anterius and anterior commissure (B and C), and brain stem (D)^13^. Arrows indicate VRS on an axially reformatted MRI slices. <u>E-F</u>: Coronally reformatted MRI slice with one nondilated VRS (E, arrow, diameter < 2 mm, dashed arrow indicates a juxtacortical MS lesion) and two dilated VRS (F, arrowhead, diameter ≥ 2 mm) from two different MS patients, both with relapsing MS phenotype. *Abbreviations: MS, multiple sclerosis; MRI, magnetic resonance imaging; VRS, Virchow-Robin spaces*.

The following predefined VRS characteristics were assessed: (1) Total VRS counts were quantified for each of above-mentioned brain levels. (2) VRS counts with a diameter of ≥ 2 mm (hereafter referred to as “dilated VRS” **Figure 1F**) were quantified on coronally reformatted T1 slices by measuring the widest diameter. The cutoff of 2 mm was based on previous studies assessing VRS^5,17,25^. (3) VRS volumes were determined by manually segmenting VRS on FLAIR-registered, coronally reformatted T1-weighted images using Freeview and FMRIB Software Library (FSL) UTILS^26^. In addition, VRS were automatically segmented using a in-house pipeline based on 3D Frangi filtering.

To investigate spatial association of VRS to MS lesions, for each subject, binary VRS and lesion masks were created. The FreeSurfer segmentation output and the LST output were used as templates for T1 and T2 lesion masks, respectively. These templates, as well as the manually segmented VRS masks, were resampled to their original T1w image using FSL’s Linear Image Registration Tool (FLIRT) and subsequently binarized using FSL UTILS^26^. In order to quantify the lesion volume in proximity to VRS, VRS masks were expanded by two voxels on each axis using FSL UTILS and the overlap between lesions and expanded VRS masks was assessed. We hypothesized that dilated VRS have a larger overlap with MS lesions compared to non-dilated VRS.

### Postmortem MRI

Postmortem MRI scans from MS brains were acquired as previously described^27^. Briefly, formalin-fixed brains were positioned in a Fomblin-filled container and were scanned in a 7-tesla MRI scanner (Siemens) equipped with a birdcage-type transmit coil and a 32-channel receive coil. A 3D T1w magnetization-prepared rapid gradient echo (T1-MP2RAGE, repetition time = 2200 ms, echo time = 3.04 ms, flip angle = 7 degrees, nominal resolution. 0.6×0.6×0.6 mm^3^, acquisition time: 6 minutes, 35 seconds) and a 3D high-resolution multigradient-echo (GRE, repetition time = 60 ms, echo times = 6.09, 15.99, 25.89, and 35.79 ms, flip angle = 10 degrees, nominal resolution. 0.42×0.42×0.42 mm^3^, acquisition time: 2 hours, 15 minutes) T2*w sequence were acquired.

### Histopathology

Histopathological validation of MRI findings was performed as described previously^28^. Briefly, brains were placed in individualized cutting-boxes and were sectioned to 1-cm-thick coronal slices. The match between the gross anatomy of the slices and the coronal T1w data was determined visually according to cortical and ventricular profiles. From these samples, VRS-corresponding tissue features were identified. We selected VRS based on their distinct macroscopic appearance on the brain slab and their correspondence to either pre-or postmortem MRI. Subsequently, these brain regions were sliced to 5-μm-thick sections using a microtome. Slides were stained with hematoxylin and eosin (H&E), Luxol fast blue-periodic acid Schiff (LFB-PAS), and, to assess vascular pathology, with Verhoeff van Gieson as well as congo red^29^. Additionally, a panel of different immunohistochemistry stains was applied to evaluate anatomy and pathology of the perivascular spaces and the adjacent CNS parenchyma, including CD3, Iba1, PLP, NFL, fibrinogen, laminin a1, collagen IV a1, and AQP-4 (**Supplementary Table 1)**.

Two raters (KBMF, a board-certified neuropathologist, and BVI) independently assessed pathological VRS features in a blinded fashion. Blood vessels were identified as veins or arteries based on Verhoeff van Gieson staining. CD3, PLP, and neurofilament light chain immune stainings were used to assess immune cellularity, (de)myelination status, and axonal damage adjacent to dilated and nondilated VRS, respectively. All stainings were semiquantitatively evaluated to assess respective pathology (score from 0-4 for each outcome). Additionally, vascular disease was semiquantitatively assessed by the presence of endothelial proliferation, splitting of the lamina elastica interna, microatheroma (i.e., presence of lymphocytic/monocytic infiltration to vessel walls), concentric hyaline thickening, perivascular hemosiderin deposition, and/or vascular tortuosity^29^ (each accounting for 1 point, i.e., score of 0–6 per vessel).

### Statistical analyses

Statistical analyses were performed using R statistical software version 3.5.2. Univariate linear regression modelling was used to assess potential associations between VRS and outcome measures which were, if explicitly stated, adjusted for sex, age, and vascular risk factors. Group differences were tested using a two-tailed t-test in case of normally distributed data and a Mann-Whitney U-test in case of nonparametric data. P values <0.05 were considered statistically significant. In order to assess the association between VRS dynamics and other imaging and clinical features, patients were stratified into groups with either VRS volume increase or decrease to retain sufficient statistical power. Multiple comparisons adjustment using a Benjamini-Hochberg correction was applied to control for false discovery rates (associaton of VRS to demographic parameters: 10 tests; comparison of VRS measures between MS and controls: 12 tests; association between VRS and MS lesions: 4 tests; proximity analysis: 1 test; pre-postmortem MRI and histopathology: 4 tests). Corrected p values are reported throughout the paper.

### Study approval

All clinical studies were approved by the regional ethics review boards (STOP-MS No. 2009/2017-31/2, last amendment 2022-01015-02; MultipleMS 2017/1323-31, amended 2018/2713-32; MyelinMS No. 2018/903-31/2), and informed consent was obtained. The formalin-fixed brains were attained at autopsy after consent was obtained from the next of kin.

## Results

### Cohort demographics and participant characteristics

Three separate clinical cohorts were included in this study: (1) the exploratory MS cohort including 142 MS patients; (2) the validation MS cohort including 63 RRMS patients; and (3) an age-matched healthy control cohort including 30 individuals. There was no difference in sex or age distribution between either of the cohorts. In total, 38 of 142 subjects (27%) had at least one vascular risk factor. At time of imaging, 171 of 205 MS patients underwent disease modifying therapy (DMT; 50 interferon beta, 50 rituximab, 32 dimethyl fumarate, 25 natalizumab, 14 glatiramer acetate) and 30 patients did not receive DMT. Participant demographics and disease characteristics are summarized in **Table 1**.

**Table 1:** Participant demographics and disease characteristics at baseline. Data shown for the two MS cohorts (exploratory and validation cohort) as well as for the control cohort. The validation cohort had a shorter disease duration compared to the exploratory cohort (p = 0.02, Mann Whitney U test). There were no statistically significant differences between the cohorts regarding sex, MS phenotype, age, or EDSS. Median estimated disease duration is based on time since symptom onset. Asterisks indicate statistical significance between cohorts: *p < 0.05. *Abbreviations: BPF, brain parenchymal fraction; EDSS, Expanded Disability Status Scale; IQR, inter-quartile range; SD, standard deviation*.

| Cohort | Exploratory | Control | Validation |
| --- | --- | --- | --- |
| **Number of subjects** | **142** | **30** | **63** |
| **Sex** |  |  |  |
| Female | 95 (67%) | 23 (77%) | 48 (76%) |
| Male | 47 (33%) | 7 (23%) | 15 (24%) |
| **MS phenotype** |  |  |  |
| Relapsing-remitting | 113 (80%) | N/A | 63 (100%) |
| Secondary-progressive | 23 (16%) | N/A | 0 |
| Relapsing-progressive | 2 (1%) | N/A | 0 |
| Primary-progressive | 4(3%) | N/A | 0 |
| **Mean age (±SD) [years]** | **37 (± 6)** | **38 (± 12)** | **36 (± 3)** |
| **Median EDSS (IQR)** | **2.0 (1.1 – 3.5)** | **N/A** | **1.0 (0.5 – 2.5)** |
| **Median disease duration (IQR) [years]** | **6.3 (1.8 – 10.6)\*** | **N/A** | **3.4 (1.2 – 6.8)\*** |
| **Number of MRI scans** |  |  |  |
| 1 scan | 142 patients | 30 subjects | 63 patients |
| 2 scans | 20 patients | 0 | 0 |
| 3 scans | 4 patients | 0 | 0 |
| **Median follow-up (IQR) [months]** | **11 (7 – 25)** | **N/A** | **N/A** |
| **Disease characteristics** |  |  |  |
| Median T1 lesion volume(±IQR) [ml] | 1.7 (± 1.2) | N/A | 0.3 (± 0.8) |
| Median T2 lesion volume (±IQR) [ml] | 2.6 (± 2.5) | N/A | 1.3 (± 0.4) |
| Median BPF (±IQR) [%] | 72 (± 2.0) | 77 (± 3.2) | 74 (± 1.4) |
| **Vascular risk factors** |  |  |  |
| Diabetes | 9 (6%) | N/A | N/A |
| Hypertension | 21 (15%) | N/A | N/A |
| Dyslipidemia | 11 (8%) | N/A | N/A |
| Antiplatelet | 11 (8%) | N/A | N/A |
| Smoking | 17 (12%) | N/A | N/A |

### Age and sex as contributor to VRS counts and volumes in the primary cohort

There was substantial agreement in Cohen’s kappa for VRS counts and volume between two raters (κ = 0.69-0.72, p<0.001).

In the exploratory MS cohort, higher age and male sex were associated with higher centrum semiovale VRS counts (β = 0.02, p = 0.02 and β = 0.31, p = 0.04, respectively) and volumes (β = 3.69 μl, p = 0.005 and β = 66.18 μl, p = 0.02, respectively). Female sex was a significant positive contributor to basal ganglia VRS (β = 0.28, p < 0.001). Increasing age was a significant positive contributor to basal ganglia VRS volume (β = 0.93, p = 0.01). There were no associations with age or sex for dilated VRS (i.e., VRS with a diameter ≥ 2 mm). These associations were not reproduced in the validation or control cohorts, hence while we adjusted for age and sex in subsequent analyses, we did not consider disease status as a modifier of these potential associations.

### MS patients have greater VRS counts and volumes compared to control subjects

Median VRS counts differed significantly between MS patients and controls in the centrum semiovale (exploratory cohort: 7 [IQR: 4 – 13]; validation cohort: 10 [6 – 16] versus control 4 [2 – 6], p = 0.002 and p < 0.001, respectively) and basal ganglia (exploratory cohort: 10 [8 – 12], validation cohort: 6 [4 – 7] versus control 4 [3 – 6], p < 0.01; **Table 2** and **Figure 2A**). Similarly, the median VRS volume (IQR) in the centrum semiovale was greater in MS patients compared to controls (exploratory cohort: 168 μl [60 – 240]; validation cohort: 263 μl [135 – 464] versus control 90 μl [50 – 181], p = 0.03 and p < 0.001, respectively; **Table 2** and **Figure 2B**). Finally, the mean count of dilated VRS was greater in MS patients compared to controls (exploratory cohort: 0.82 [±SD: 0.69]; validation cohort: 1.21 [± 1.63] versus control 0.2 [± 0.61], p = 0.04 and p = 0.002, respectively, **Table 2** and **Figure 2C**).

**Table 2:** Virchow-Robin space (VRS) prevalence, counts, and volumes per location for the two MS cohorts. Higher centrum semiovale and basal ganglia VRS counts and centrum semiovale VRS volumes in MS patients compared to controls (Exploratory cohort: n = 142; validation cohort: n = 63; control cohort n = 30). Asterisks indicate statistical significance between cohorts: *p < 0.05, **p < 0.01, ***p < 0.001. *Abbreviations: IQR, interquartile range; N/A, not available; SD, standard deviation*.

| Cohort | Exploratory | Control | Validation |
| --- | --- | --- | --- |
| Number of subjects | 142 | 30 | 63 |
| **Prevalence of VRS per location (%)** |  |  |  |
| Centrum semiovale | 137/142 (96%) | 30/30 (100%) | 63/63 (100%) |
| Basal ganglia | 142/142 (100%) | 30/30 (100%) | 63/63 (100%) |
| Brain stem | 44/142 (31%) | 14/30 (47%) | 15/63 (24%) |
| **Median VRS count per location (IQR)** |  |  |  |
| Centrum semiovale | 7 (4 – 13) ** | 4 (2 – 6) *** | 10 (6 – 16) |
| Basal ganglia | 10 (8 – 12) ** | 4 (3 – 6) | 6 (4 – 7) |
| Brain stem | 0 (0 – 1) | 0 (0 – 1) | 0 (0 – 0) |
| **Median VRS volume per location [μl] (IQR)** |  |  |  |
| Centrum semiovale | 168 (60 – 240) * | 90 (50 – 181) *** | 263 (135 -464) |
| Basal ganglia | 136 (112 – 169) | 158 (128 – 208) | 146 (114 -176) |
| Brain stem | N/A | N/A | N/A |
| **Mean count of dilated VRS (diameter ≥ 2 mm, ± SD)** |  |  |  |
| Centrum semiovale | 0.82 (± 1.69) * | 0.2 (± 0.61) ** | 1.21 (± 1.63) |
| No of Patients with VRS ≥ 2 mm (%) | 43 (30%) | 4 (13%) | 31 (49%) |

**Figure 2:**
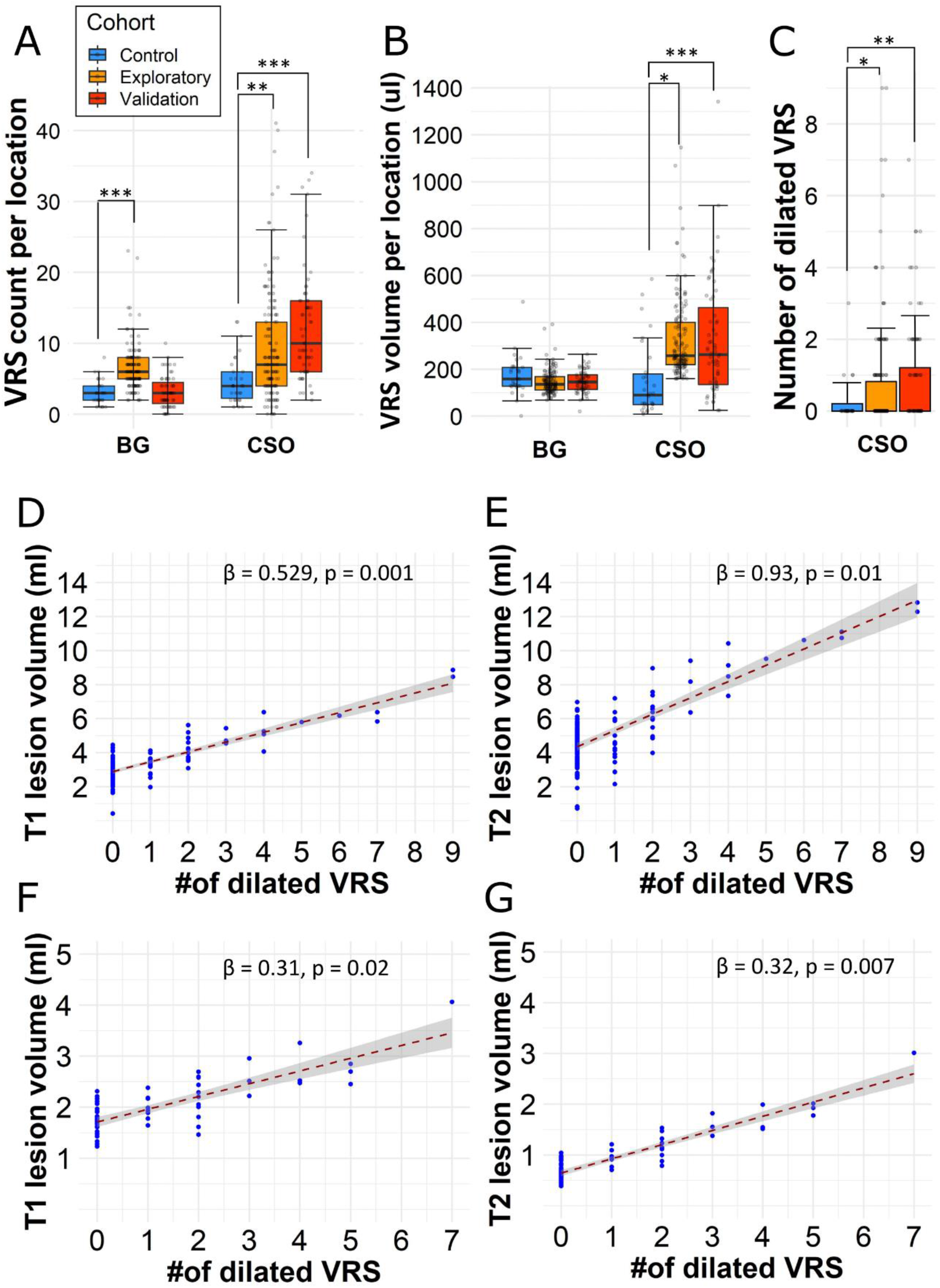
Dilated VRS are associated with higher T1 and T2 lesion volumes. Higher VRS counts (A), volumes (B), and number of dilated VRS (diameter ≥ 2 mm, C) in MS patients compared to controls (Exploratory cohort: n = 142; validation cohort: n = 63; control cohort: n = 30). The count of dilated perivascular spaces (VRS, diameter ≥ 2 mm) was associated with higher MRI (D) T1 lesion and T2 lesion volume in our exploratory cohort (n=142). These associations were substantiated in the validation cohort (n=63, F and G). The corresponding slope of the regression line (β) and p values are displayed at the bottom right corner of each graph. The regression models are adjusted for patient age and sex. Asterisks indicate statistical significance between cohorts: *p < 0.05, **p < 0.01, ***p < 0.001. *Abbreviations: BG(A/C), basal ganglia (anterior commissure/crus anterius level); CSO, centrum semi-ovale; VRS, Virchow-Robin space*.

### The number of dilated VRS is associated with T1 and T2 lesion volumes

VRS counts and volumes were not associated with T1 or T2 lesion volume. Interestingly, however, the number of dilated VRS (diameter ≥ 2 mm) in the centrum semiovale was associated with both T1 and T2 lesion volumes (β = 0.53 ml, p < 0.001 and β = 0.93 ml, p = 0.01, respectively). These findings remained statistically significant in multivariable analysis with age, sex, supratentorial brain volume, and number of vascular risk factors as independent variables (T1 lesions: β = 0.53, p = 0.001 and T2 lesions: β = 0.93, p = 0.01; **Figure 2D and E**). Additionally, findings were corroborated in the validation cohort, in which we also found associations between the number of dilated VRS and T1 or T2 lesion volumes (β = 0.31, p = 0.02 and β = 0.32, p = 0.007, respectively; **Figure 2F and G**). As expected, T1 and T2 lesion volume were positively correlated to each other (r = 0.88, p < 0.001). No such association was found for basal ganglia VRS in either of the cohorts. No association was found between VRS counts, volume or number of dilated VRS and volume of Gd-enhancing lesions. VRS counts, volumes, or diameters were not associated with brain parenchymal fraction in the uni-or multivariable analysis (data not shown).

### No association of VRS with clinical parameters

We did not find an association of VRS counts, diameters, or volumes with EDSS or count of total relapses. We also did not find an association of VRS counts or volumes with cognitive performance of MS patients as assessed by SDMT.

### No bias of T1 or T2 lesions to dilated VRS compared to non-dilated VRS

Both dilated and non-dilated centrum semiovale VRS were mainly located at the parieto-occipital transition region and in the superior cerebral convexities (**Figure 3A – C**). Basal ganglia VRS were mainly located in the dorsal aspects of the putamen and the globus pallidus (**Figure 3D – F**). Control subjects had a similar dispersion of centrum semiovale and basal ganglia VRS (data not shown).

**Figure 3:**
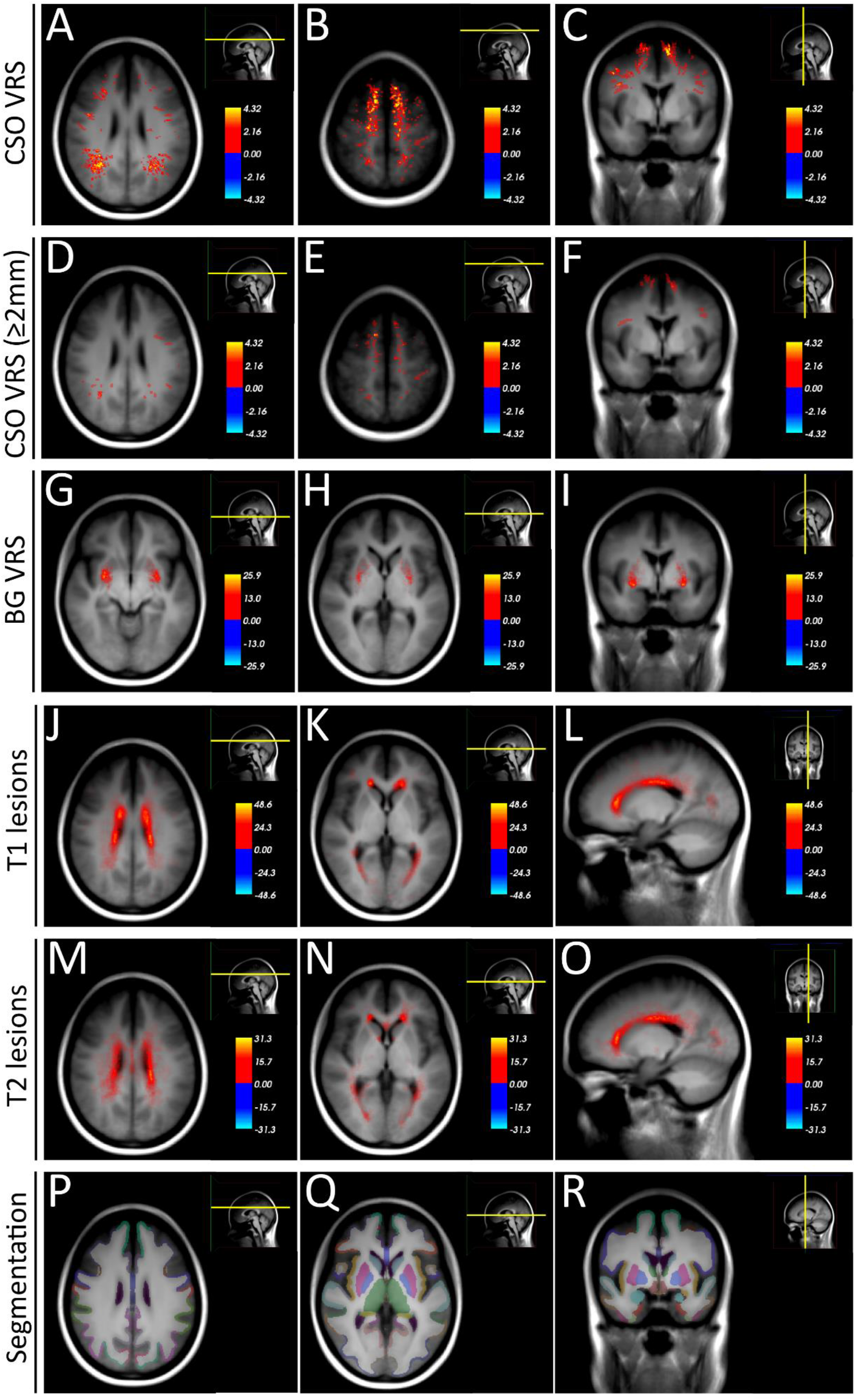
VRS and MS lesion heat maps in the exploratory cohort. Heat maps of VRS and T1 lesions in the exploratory MS cohort (n = 142), superimposed on the average space MRI scan of the exploratory cohort. VRS are mainly located at the parieto-occipital transition region and in the superior convexities (A – C). Dilated VRS (diameter ≥ 2 mm) show a similar distribution (D – F). Basal ganglia VRS are mainly located in the dorsal aspects of the putamen and the globus pallidus (G – I). T1 and T2 lesions are mainly located adjacent to the lateral ventricles (J – O). For comparison, P – R displays the cortical and subcortical parcellation/segmentation output from Free-Surfer^47^. *Abbreviations: BG, basal ganglia; CSO, centrum semiovale; MS, multiple sclerosis, VRS; Virchow-Robin spaces*.

There was only very small overlap of VRS with MS lesions **(Figure 4)**. There was no statistically significant difference in proximity of either T1 or T2 lesions to dilated or non-dilated centrum semiovale VRS in the validation cohort (Wilcoxon signed-rank test, for T1 lesions p = 0.33; for T2 lesions p = 0.21, **Figure 4B**).

**Figure 4:**
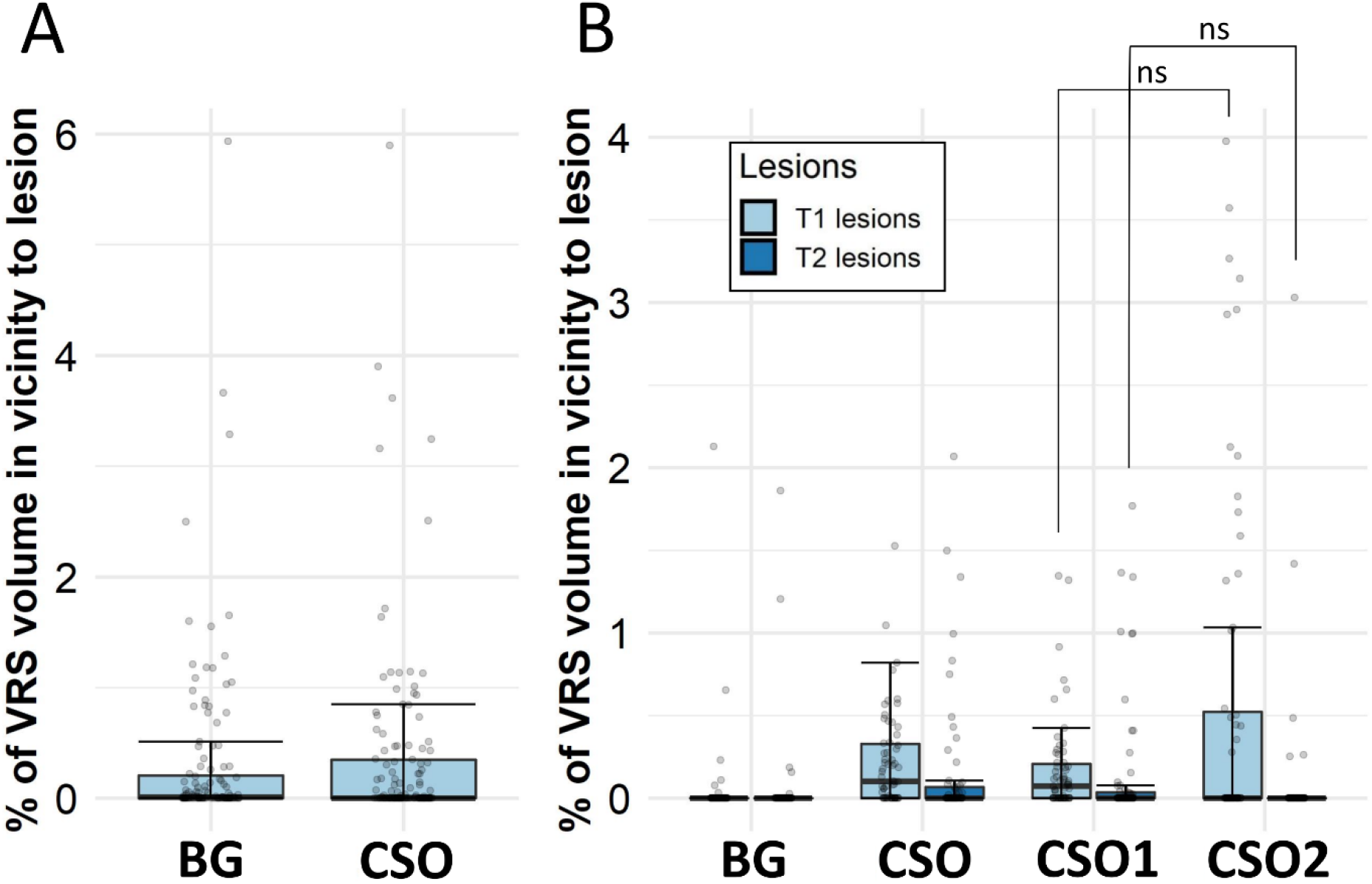
Quantification of overlap between VRS and MS lesions. Percentage of volume overlap between Virchow-Robin spaces (VRS, dilated by two voxels on each side) and MS lesions in (A) the exploratory (n = 142) and (B) the validation multiple sclerosis (MS) cohort (n = 63). For the exploratory cohort, only VRS proximity to T1 lesions is displayed. Many patients show zero overlap, with some cases showing up to 6% overlap between VRS and MS lesions. There is no statistically significant difference between nondilated (CSO1) and dilated (CSO2) centrum semiovale VRS in overlap with either T1 or T2 lesions. In (B): Left box/whiskers of the corresponding VRS location represents proximity to T1 lesions and right box/whiskers T2 lesions. *Abbreviations: BG, basal ganglia; SOC, centrum semiovale; VRS, Virchow-Robin spaces*.

### VRS show restrained temporal evolution

Our exploratory cohort included 24 patients with up to 3 longitudinal MRI scans and a cumulative follow-up time of 31.5 patient-years (median follow-up time 18 months [IQR: 12 – 24]). During the followup, 5 patients presented with at least 1 new T2 lesions during the follow-up and 2 patients changed DMT (from betaferon to rituximab). During the follow-up period, a total of 30 new VRS were detected, corresponding to around 1 new VRS per patient-year. Most new VRS were detected in the basal ganglia (0.51/year), followed by the centrum semiovale (0.30/year).

Overall, VRS volumes did not change significantly during the observation period in either the basal ganglia or the centrum semiovale (**Supplementary Table 2**). Regarding evolution of VRS diameters: 12 out of 24 longitudinally followed-up patients showed a total of 35 dilated VRS at the initial scans. Seven of these 12 patients (58%) showed a total increase of 10 dilated VRS within 31.5 patient years. Finally, there was no association of VRS volume change with baseline or longitudinal imaging (gadolinium enhancing lesion volume, T1 or T2 lesion volume as well as brain volume/atrophy measures) or clinical outcome measures (number of total relapses, EDSS, SDMT).

### Pre-postmortem dynamics of VRS

Our postmortem cohort comprised 6 MS patients (5 with a progressive, 1 with a relapsing clinical phenotype, **Table 3**). First, we assessed interrater agreement for VRS detection on postmortem MRI: there was substantial agreement in Cohen’s kappa (κ = 0.61-0.71, p<0.001). Next, we assessed whether VRS appearance in MRI changed upon death in 3 of the patients with both pre- and postmortem MRI. Qualitatively, although all VRS were identifiable on both pre- and postmortem MRI, VRS appeared more distinct on T1w premortem scans compared to postmortem scans. This was confirmed in a quantitative analysis measuring VRS volume: Mean VRS volume was 12.6 μl (±SD 10.2) in premortem scans and 1.8 μl (±1.2) on postmortem scans (9 VRS, p=0.006). In contrast, MS lesion volumes did not change significantly pre-versus postmortem (premortem: 63.6±47.7 μl, postmortem: 53.4±48.1 μl, 8 MS lesions, p=0.68) (**Figure 5A-C**).

**Table 3:** Postmortem multiple sclerosis (MS) cohort. Brains from 6 multiple sclerosis (MS) patients and 3 control subjects were investigated for histopathological validation of (dilated) Virchow-Robin spaces (VRS). *Abbreviations: MRI, magnetic resonance imaging; PPMS, primary progressive multiple sclerosis; SPMS, secondary progressive multiple sclerosis; RRMS, relapsing-remitting multiple sclerosis. *No postmortem MRI available. The VRS burden was evaluated macroscopically on brain slabs*.

| ID, sex, age | Diagnosis | Cause of death | Postmortem interval | MRI VRS burden | Number of retrieved tissue blocks |
| --- | --- | --- | --- | --- | --- |
| **MS patients** |  |  |  |  |  |
| MS1, M, 60 | PPMS | Brainstem stroke | 9 hours | High | 14 |
| MS2, F, 60 | Progressive MS | Sepsis | 7.5 hours | Medium to high | 5 |
| MS3, F, 78 | Progressive MS | Pneumonia | 26 hours | Medium | 7 |
| MS4, F, 57 | SPMS | Unknown | 89 hours | High* | 5 |
| MS5, F, 78 | Progressive MS | Unknown | 12-15 hours | Medium | 2 |
| MS6, F, 76 | RRMS | Unknown | <24 hours | High | 9 |
| **Controls** |  |  |  |  |  |
| C1, M, 46 | X-ALD | Sepsis | 48 hours | Low | 4 |
| C2, F, 71 | Brain metastases (primary tumour unknown) | Unknown | <24 hours | Medium | 3 |
| C3, M, 62 | Subarachnoid bleeding | Pneumonia | <24 hours | Medium | 2 |

**Figure 5:**
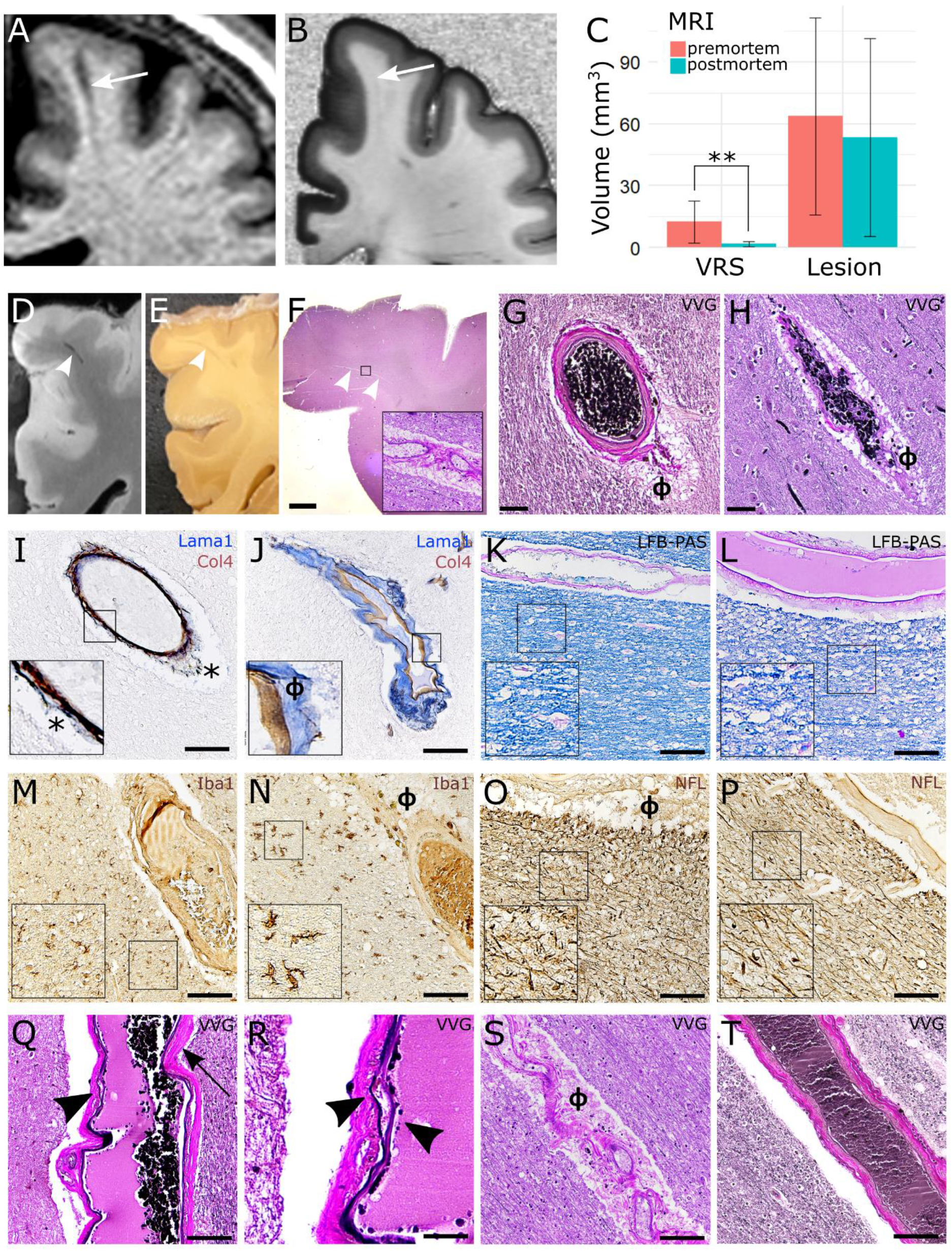
Histopathological validation of dilated and non-dilated VRS. <u>A-C</u>: Dynamic changes of pre-(A) versus postmortem VRS (B): VRS show a substantial drop in volume from pre-to postmortem MRI (quantification: C). <u>D-F</u>: VRS, as identified on postmortem T1w MRI scans (D), were correlated to corresponding brain tissue blocks I and paraffin tissue slices (F, inlet with 200x magnification). VRS corresponded mostly to arteries (G, φ = perivascular space) and to a lesser extent veins (H). <u>I and J</u>: The perivascular compartments were identified with double staining for laminin α1 (labelling the parenchymal basement membrane, blue) and collagen IV α1 (labelling the vascular basement membrane, brown) (J) which was in contrast to artifacts caused by perimortem vascular collapse (asterisk, I; inlets with 200x magnification). <u>K-P</u>: Neither dilated (K, M, O, Q, S) nor nondilated VRS (L, N, P, R, T) were associated with demyelination (K and L, LFB-PAS), activation of micro-glia/macrophages (M and N, Iba1), or axonal damage (O and P, neurofilament [NFL]; insets with 200x magnification]. <u>Q-T</u>: Dilated VRS of the arterial tree were more commonly associated with signs of small vessel disease, such as splitting of the internal elastic lamina (arrowhead in Q, 200x magnification, higher magnification in R), vessel wall hyalinosis (arrow in Q), or vascular tortuosity (S) compared to nondilated VRS (T). Scale bars = 50 μm, except in F = 1mm and R = 10 μm. *Abbreviations: φ, perivascular space; LFB-PAS, Luxol Fast Blue periodic acid-Schiff; MRI, magnetic resonance imaging; VVG, Verhoeff van Gieson; VRS, Virchow-Robin space*.

### Histopathological validation of VRS

To assess association of VRS with veins or arteries, VRS were localized on postmortem T1w MRI scans and subsequently correlated to their corresponding tissue substrate (**Figure 5D-F**). We assessed 19 VRS in MS cases and 6 VRS in control subjects. In the centrum semiovale, most VRS were associated with arterial vessels: out of 19 assessed VRS, 16 were associated with arteries (84%, **Figure 5G**) and 3 with veins (**Figure 5H**). In the basal ganglia, all VRS were associated with arteries (7 VRS, data not shown). Of note, perimortem collapse of blood vessels can cause artificial widening of the perivascular space. Hence, we additionally identified the true perivascular compartment with a double staining for laminin α1 (labelling the parenchymal basement membrane) and collagen IV α1 (labelling the vascular basement membrane) (**Figure 5I and J**).

Finally, we assessed the potential association of VRS with MS pathology features. Neither dilated nor non-dilated VRS were associated with adjacent demyelination (dilated VRS score: 0.75[±SD: 0.71], non-dilated VRS score: 0.96[±SD: 0.71]), perivascular cuffs (1[±SD: 1.14], 0.81[±SD: 1.01]), macrophage/microglial activation (2[±SD: 2.14], 0.82[±SD: 0.69]), or axonal damage (number of axons: 204[±SD: 24], 211[±SD: 78]) (**Figure 5K-P**). Furthermore, VRS were not associated with perivascular amyloid deposition (data not shown). VRS were also not associated with perivascular fibrin deposition indicating blood-brain barrier (BBB) leakage (data not shown). Interestingly, dilated VRS in the centrum semiovale were more commonly associated with signs of small vessel disease, such as splitting of the internal elastic lamina or vessel wall hyalinosis (**Figure 5Q-S**), compared to non-dilated VRS (**Figure 5T**). This was confirmed in a quantitative analysis comparing the small vessel disease scores in dilated versus non-dilated VRS (dilated VRS: 3.86[±SD: 1.34], non-dilated VRS: 2.21[±SD: 0.89], p=0.03). Such an association was not found in 3 control subjects without MS (6 VRS, data not shown).

## Discussion

### Main findings

The objective of this study was to investigate the potential association of nondilated and dilated VRS with clinical and imaging parameters in MS. Furthermore, we assessed the histopathological signature of VRS in a postmortem cohort of 6 MS patients. In the MS cohorts, the count of dilated centrum semiovale VRS (diameter ≥ 2 mm), but not total VRS counts or volumes, was associated with increased T1 and T2 lesion volumes. VRS did not colocalize with T1 and T2 lesions at the anatomical level. Instead, they mostly corresponded to arteries and were not associated with common MS pathology features such as demyelination, axonal damage, or immune cell infiltration. However, intriguingly, the tissue signature of dilated VRS in the centrum semiovale corresponded to signs of arterial disease, despite the fact that focal demyelination in the MS white matter is a perivenular process.

### Findings in the context of existing evidence

We have recently conducted a meta-analysis that substantiated the notion that MS patients carry a higher VRS burden compared to controls^15^. Findings from our current original study are in line with this notion. Additionally, in the current study, the count of dilated centrum semiovale VRS (diameter ≥ 2 mm), but not their volume or count, was associated with increased T1 and T2 lesion volume. Likewise, two other previous studies did not find an association of total VRS volume and/or count with T1 and/or T2 lesion burden^14,16^. However, these two studies did not assess dilated VRS. In contrast to the association with T1 and T2 lesion volumes, our analyses did not disclose an association between VRS measures and physical or cognitive disability in MS. It might be that our analyses were confounded by the relatively low level of physical and cognitive disability in our cohorts.

The association of VRS with neuroinflammatory pathology has been shown previously. Notably, a longitudinal study observed a VRS volume increase preceding the emergence of contrast-enhancing MS lesions^16^ Thereupon, it has been speculated that the antecedent surge in VRS volume could represent a local accumulation of immune cells within the perivascular spaces. Focal perivascular space dilation has also been observed in a case series at the edges of active MS lesions at the initiation of inflammatory exacerbation^30^ However, in our longitudinal cohort, we did not find an association between VRS outcomes and gadolinium-enhancing, T1, or T2 lesions. Furthermore, T1 and T2 lesions did not spatially cluster with VRS. Also, the absence of common MS pathology features adjacent to VRS, including the lack of perivascular cuffing, renders their direct association with neuroinflammatory tissue pathology unlikely.

Clinical MRI at conventional field strengths cannot easily discriminate perforating arterioles and venules. Thus, it is currently under debate whether VRS surround arterial or venous vessels or both^2,7^. A small imaging study employing structural T2w MRI scans in conjunction with angiographies at ultra-high static magnetic field strengths found that VRS correspond to arterial vessels^31^. Our MRI-histology correlation of VRS corroborates this observation by providing histological evidence that VRS in the centrum semiovale and basal ganglia mostly correspond to arteries. Because perivascular cuffing in MS is commonly observed surrounding venules^32^, this notion further argues against the hypothesis that dilation of VRS corresponds to perivascular cuffing^16^.

An increase of VRS has been consistently associated with ageing^8^, which is also confirmed by our data from MS patients, though naturally chronologically confounded by the disease duration. Based on this association with ageing, it has also been speculated that VRS represent a perivascular *ex vacuo* atrophy ^3^. In MS, a study employing ultra-high-field MRI found that higher VRS counts were associated with lower brain parenchymal fraction^13^. Our study does not support this observation: dilation of VRS was not explained by higher degrees of brain volume loss in our MS cohort. Furthermore, we did not find myelin pallor or apparent loss of axonal density as signs of Wallerian degeneration adjacent to VRS^33^. Nevertheless, more widespread neurodegeneration and/or volume loss of extra-neuronal tissue components cannot easily be ruled out by histopathological analysis.

VRS have been associated with small vessel disease^6^ and cardiovascular risk factors such as hypertension^34^. Interestingly, in cerebral amyloid angiopathy, the degree of VRS dilation was associated with more pronounced accumulation of Aβ in upstream juxtacortical vessel walls^35^. While we did not observe vascular Aβ deposition in VRS in our histopathological analysis, we did find more distinct signs of arterial pathology in dilated versus nondilated VRS in MS compared to non-MS controls. This is in line with observations from a large postmortem study in MS patients reporting higher cerebral small vessel disease features such as periarteriolar space dilation in MS patients^36^. Interestingly, vascular comorbidities have been associated with worse cognitive function^37^ and lower brain volumes in MS^38^ (reviewed in^39^). This is in line with our observations that dilated VRS — corresponding to arteries with vascular disease — are associated with higher T1 and T2 lesion volumes. This opens the question about the pathological cascade of these events, i.e., does vascular disease precipitate neuroinflammation or does chronic neuroinflammation and its downstream effects cause vascular disease? A large genome-wide association study did not provide evidence to suggest a shared genetic mechanism of ischemic white matter damage and MS^40^. Furthermore, the association between dilated VRS and lesion volumes was statistically independent from general vascular risk factors in our study, which may indicate a more direct causal link between VRS and MS pathology. In contrast, it has been shown in a neuroinflammatory marmoset model that intralesional veins can show vascular remodeling with perivascular collagen deposition already during early lesion emergence^41^. It can be speculated that similar mechanisms might contribute to arterial pathology in chronic and widespread neuroinflammation like in MS.

There is insufficient understanding of how VRS become dilated^5^: it has been hypothesized that perivascular fibrosis, *ex vacuo* atrophy of adjacent brain tissue, or alterations of arterial wall permeability might cause dilation. Our data add the notion that arterial disease could also cause VRS dilation. In addition, the local widening of the perivascular spaces in the juxtacortical white matter, e.g. in the centrum semiovale, could indicate impaired interstitial or cerebrospinal fluid drainage^42^ and/or excess fluid leakage from the vasculature^43^. In our pre- and postmortem MRI comparison, the numbers of VRS remained similar, in line with findings from a recent MRI-postmortem study^44^. However, we observed a strong decrease in VRS volumes upon death. It is unclear to what extent this comes from a drop of blood or CSF pressure or both, but this observation could allude toward increased local CSF pressure. To further test this hypothesis, sensitive imaging methods are warranted to visualize BBB pathology and/or perivascular fluid drainage in proximity to VRS, such as dynamic contrast enhanced MRI^45^ or CSF tracer studies^46^.

### Limitations

Our study has some limitations: (1) Our longitudinal cohort only included 21 patients with longitudinal scans (followed-up for 31.5 patient years). Although this is one of the largest longitudinal cohorts of VRS in MS to date, the cohort size is relatively small compared with those investigated for other MRI outcomes. Development of automated tools for VRS identification and segmentation could facilitate larger studies with longer follow-up. (2) In our postmortem cohort, 5 of 6 patients had a progressive clinical MS phenotype, contrasting with the mostly relapsing clinical phenotype of the *in vivo* cohort. This predominance of more chronic disease could have biased our analysis.

### Conclusions

In MS, higher numbers of centrum semiovale dilated VRS were associated with greater T1 and T2 lesion volumes. Correlative histopathology showed that these VRS mostly corresponded to arteries displaying signs of vascular disease but no MS pathology hallmarks. Together, these data argue against the hypothesis that the dilation of perivascular spaces represents a surge of immune cells prior to a neuroinflammatory outbreak, but instead point towards an association of MS with arterial disease. To corroborate these findings, future studies should longitudinally assess VRS dynamics and their association with markers of small vessel disease.

## Glossary

BBB: blood-brain barrier
EDSS: Expanded Disability Status Scale
EPVS: enlarged perivascular space
IQR: interquartile range
LST: lesion segmentation tool
MRI: magnetic resonance imaging
MS: multiple sclerosis
RR: relapsing-remitting
PP: primary progressive
SP: secondary progressive)
VRS: Virchow-Robin spaces

## Acknowledgments

We thank R. Schumann and J. Vandroogenbroeck for help with data analysis.

## Funding

This work was supported by grants of the Swiss National Science Foundation (No. P400PM_183884, to BVI), the UZH Alumni (to BVI), the Swedish Society for Medical Research (No. S19-0227 to TG), and by the Intramural Research Program of NINDS, NIH, USA. We thank all our funders for their support.

The sponsors had no role in the design and conduct of the study; collection, management, analysis, and interpretation of the data; preparation, review, or approval of the manuscript; and decision to submit the manuscript for publication.

## Competing interests

The authors report no competing interests.

## Supplementary material

Supplementary material is available at Brain online.

## Author contributions

Conception and design of study: BVI, TM, FP, DSR, TG; acquisition of data: BVI, MP, RO, IK, TG; analysis of data: BVI, CC, KBMF; drafting the initial manuscript: BVI; all authors critically revised the paper draft.

## Supplementary Tables

**Supplementary Table 1:** Staining protocols for immunohistochemistry panel. Immunohistochemistry staining protocols for used antigens on paraffin-embedded 5 μm thick tissue sections. *Abbreviations: BM, basement membrane; HIER, GFAP, glial fibrillary acidic protein; Heat-induced epitope retrieval; PLP, protolipid protein*.

| Antigen | Target | Vendor | Retrieval method | Concentration, incubation time |
| --- | --- | --- | --- | --- |
| PLP | Myelin | Bio-Rad | HIER with citrate buffer (pH6) | 1:200, 1h |
| Fibrinogen | Blood-brain barrier | Abcam | Proteinase K 1:1000 | 1:100, 1h |
| Collagen IV α1 | Vascular BM | Thermo Scientific (MS-747-S) | Pepsin, 37°C | 1:50, 1h |
| Laminin α1 | Glial BM | Sigma (L9393) | Pepsin, 37°C | 1:50, 1h |
| Iba-1 | Microglia/macrophages | Wako | Proteinase K | 1:400, 1h |
| CD-68 | Microglia/macrophages | Thermo Scientific | HIER with EDTA buffer (pH9) | 1:100, 1h |
| CD-3 | T Cells | Dako | HIER with EDTA buffer (pH9) | 1:100, 1h |
| GFAP | Astrocytes | Dako | Proteinase K | 1:200, 1h |
| SMI-31 | Axons | Biolegend | HIER with IHC Target Solution | 1:200, 1h |

**Supplementary Table 2:** Temporal evolution of VRS. VRS volume changes over time in 24 patients (20 patients with 2 scans, 4 patients with 3 scans) from the primary cohort. Whereas there are small changes in the overall basal ganglia VRS volume, semioval center VRS volume remained around the same level during our observation period (31.5 patient years). *Abbreviations: IQR, interquartile range; VRS, Virchow-Robin spaces*.

|  | First scan | Second scan | Third scan |
| --- | --- | --- | --- |
| Number of patients | 24 | 24 | 4 |
| VRS volume |  |  |  |
| Semioval center median (IQR) | 134 mm<sup>3</sup>(53 – 254) | 142 mm<sup>3</sup> (74 – 234) | 134 mm<sup>3</sup>(118 – 211) |
| Basal ganglia median (IQR) | 121 mm<sup>3</sup>(104 – 151) | 139 mm<sup>3</sup>(111 – 131) | 132 mm<sup>3</sup>(122 – 177) |

## Notes

**Conflict of interest statement**, The authors have declared that no conflict of interest exists.

### Competing Interest Statement

The authors have declared no competing interest.

